# Precise detection of Acrs in prokaryotes using only six features

**DOI:** 10.1101/2020.05.23.112011

**Authors:** Chuan Dong, Dong-Kai Pu, Cong Ma, Xin Wang, Qing-Feng Wen, Zhi Zeng, Feng-Biao Guo

## Abstract

Anti-CRISPR proteins (Acrs) can suppress the activity of CRISPR-Cas systems. Some viruses depend on Acrs to expand their genetic materials into the host genome which can promote species diversity. Therefore, the identification and determination of Acrs are of vital importance. In this work we developed a random forest tree-based tool, AcrDetector, to identify Acrs in the whole genomescale using merely six features. AcrDetector can achieve a mean accuracy of 99.65%, a mean recall of 75.84%, a mean precision of 99.24% and a mean F1 score of 85.97%; in multi-round, 5-fold cross-validation (30 different random states). To demonstrate that AcrDetector can identify real Acrs precisely at the whole genome-scale we performed a cross-species validation which resulted in 71.43% of real Acrs being ranked in the top 10. We applied AcrDetector to detect Acrs in the latest data. It can accurately identify 3 Acrs, which have previously been verified experimentally. A standalone version of AcrDetector is available at https://github.com/RiversDong/AcrDetector. Additionally, our result showed that most of the Acrs are transferred into their host genomes in a recent stage rather than early.

## INTRODUCTION

Prokaryotes have a variety of sophisticated and powerful adaptive defence weapons, such as R-M (restriction-modification), Abi (abortive infection), CRISPR-Cas (clustered regularly interspaced short palindromic repeats–CRISPR-associated proteins) systems (1), which can block, abolish, and stop the infection of MGEs (mobile genetic elements). Recently, Doron *et*. *al*. discovered ten immune systems, which can suppress MGEs: nine out of the ten have anti-phage functions, and the remaining one can curb plasmids (2). These immune systems constitute the pan-immune systems of prokaryotes (1). The long-term arm race between bacteria and virus has promoted the evolution of countermeasures against injury from bacterial distinct defence arsenal such as the three defence systems mentioned in the first sentence. In 2013, Bondy-Denomy *et*. *al*. reported five distinct proteins that can help bacteriophage escape immune weapons in *Pseudomonas aeruginosa* (3). They are kinds of MGEs-derived protein that disable the activity of CRISPR-Cas systems, which are known as Acrs. Some Acrs can bind with Cas proteins in CRISPR-Cas system to exert their inhibitory effect. For example, AcrIIA4 can bind with SpyCas9 to obscure the protospacer adjacent motif (PAM) interaction position (4–6). Some studies showed that some other Acrs can act as enzymes to inhibit CRISPR-Cas systems. For example, AcrVA5, as an acetyltransferase, can modify Cas12a by acetylation, thus to disable its function (7). Additionally, AcrVA1 acts as a nuclease, which can trigger the cleavage of the Cas12a-bound guide RNA to reduce the activity of Cas12a complex (8).

In the case of Acrs identification, it is difficult to pick out such proteins from a huge sequence pool because of the sequence diversity of Acrs. However, a very conserved protein, anti-CRIPSR associated protein (Aca), usually flanks downstream of Acrs. Sequence alignment has shown Aca contains HTH (helix-turn-helix) domain. The HTH domain later is utilized as a marker to search novel potential Acrs. Recent studies showed that HTH might be evolutionarily related with the toxin-antitoxin systems (9). HTH can inhibit the transcription of *acr* genes (10,11). In addition to the guilt-by-association method, comparative genomics and self-targeting (ST) rule have already used to screen candidate species that may have Acrs (3,12–15). During our manuscript preparation, some related works have posted in bioRxiv and online. For example, an available server, AcrRank (16), is available recently. More recently, an online resource, Acrs catalog, is released (17). It stores some potential Acrs and can be used as a wonderful resource to screen Acrs. Another two online servers, AcrFinder (18) and PaCRISPR (http://pacrispr.erc.monash.edu/index.jsp) were also online. Herein, we designed another tool, AcrDetector to screen Acrs in the whole genome-scale. Compared with AcrRank, which used 412 features, AcrDetector only adopts 6 features. Our designed AcrDetector can be used as a supplementary tool for the above resources. AcrDetector considers the features from the whole genome panoramic picture, whereas AcrRank leverages local features derived from sequence itself. Therefore, the they are complementary in terms of methodology.

## MATERIAL AND METHOD

### Benchmark construction

To train a model, a high-quality training dataset is necessary. We downloaded Acrs from anti-CRISPRdb (19), then carried out PSI-BLAST search against prokaryotic genomes (20) with an e-value of 10e-4. The prokaryotic genomes were downloaded from NCBI in November, 2018. We then employed rigorous thresholds (identity greater than 40% and coverage greater than 75%) to further filter homologous sequences if the sequences have clear and definite function annotation, otherwise sequences without clear and definite function annotation such as “Hypothetical” another threshold (identity larger than 40) was employed. There are ~8,363,664 sequences in the species that contained Acr homologs. Faced with such huge data, we based on downsampling method to reduce the data scale via uniform sampling on each chromosome of each species using an interval of 30 genes after excluding all the Acr homologs. The sampled sequences were regarded as negatives. On one hand the sampling method can give a curtailment of negatives, on the other hand part of sequences in each species are included, thus covering all species containing Acr homologs. Due to our method considered features of neighbours of a target gene, we also removed some genes that do not have continuous gene neighbours. Finally, a benchmark dataset including 4,131 positives and 290,498 negatives was constructed. 5 out of the 6 features employed by us depend on the genome background itself instead of features deriving from sequence (apart from the length of a target sequence). The homologs should be covered as more species as possible so that we can capture more comprehensive features for each Acrs in different genomic background. Considering the above, we do not remove redundant sequences.

### Feature extraction

Herein, we employed 6 features (Table 1) to describe entries in our benchmark. They are gene length, gene location on chromosome strand (forward and reverse strand), conserved domain in the downstream, codon usage against the whole genome, codon usage deviation compared with the whole genome-scale, and functional annotation, respectively. Gene length is defined as the nucleotides number in a sequence. We considered gene length of three genes: a target gene, left neighbour of a target gene and right neighbour of a target gene so that three variables were derived from gene length. We used binary variables to represent a target gene’s function, in which “1” means no validated function, such as “hypothetical protein” and “0” means there is a clear and definite function NCBI. Gene location on chromosome strand refers to whether a target gene and one of its two neighbours (left neighbour and right neighbour) are in the same strand. “1” was conferred to a target gene if the target gene and one of its neighbours are in the same DNA strand, otherwise “0” was conferred to a target gene. Conserved domain refers to whether there is an HTH domain in the downstream of a target gene. To search such domain, we merely considered three downstream genes of a target gene because 26 out of 45 validated Acrs have an HTH domain within three downstream genes, which is illustrated by Figure 1 in reference (9). Codon usage against the whole genome-scale is calculated by the following equation (A). Where *f*_*j*_ denotes the *j*^*th*^ codon frequency in a certain gene, and *f*_*wj*_ represents the *j*^*th*^ codon frequency in the whole genome-scale. Therefore, *D* can be utilized to estimate codon usage deviation. For each gene in a genome, we can calculate *D* value, which allows us to perform a comparison of any two genes. To further measure *D*, we proposed *dev* parameter, which can be calculated by the following equation (B). Where *I*(*D*_*i*_ > *D*_*j*_) means a comparison between *D*_*i*_ and *D*_*j*_ of gene *i* and gene *j*. The n in equation (B) represents protein coding gene number in a genome.

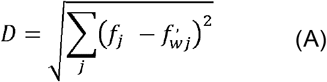

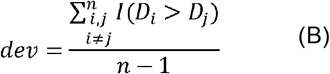

**Table 1.** Feature summary. Feature: feature name; Num: variable number derived from the corresponding feature.

| Feature | Num | Description |
| --- | --- | --- |
| Gene length | 3 | Length of left neighbour, right neighbour of a target gene plus a target gene length, therefore totally 3 variables derived from gene length. |
| Gene location on chromosome strand | 1 | A binary value to measure whether the target and one of its neighbours are in the same strand |
| Conserved domain in downstream of a target gene | 1 | A binary value to represent whether there is an HTH domain in the downstream. |
| Codon usage against the whole genome | 1 | A float value calculated by equation (A) |
| Codon usage deviation compared with the whole-genome | 3 | Float values of left neighbour, right neighbour of a target gene plus a target gene. It can be calculated by equation (B) |
| Function | 1 | A binary value to represent the function. "1" unknown function. "0" a validated function in NCBI annotation |

**Figure 1a.**
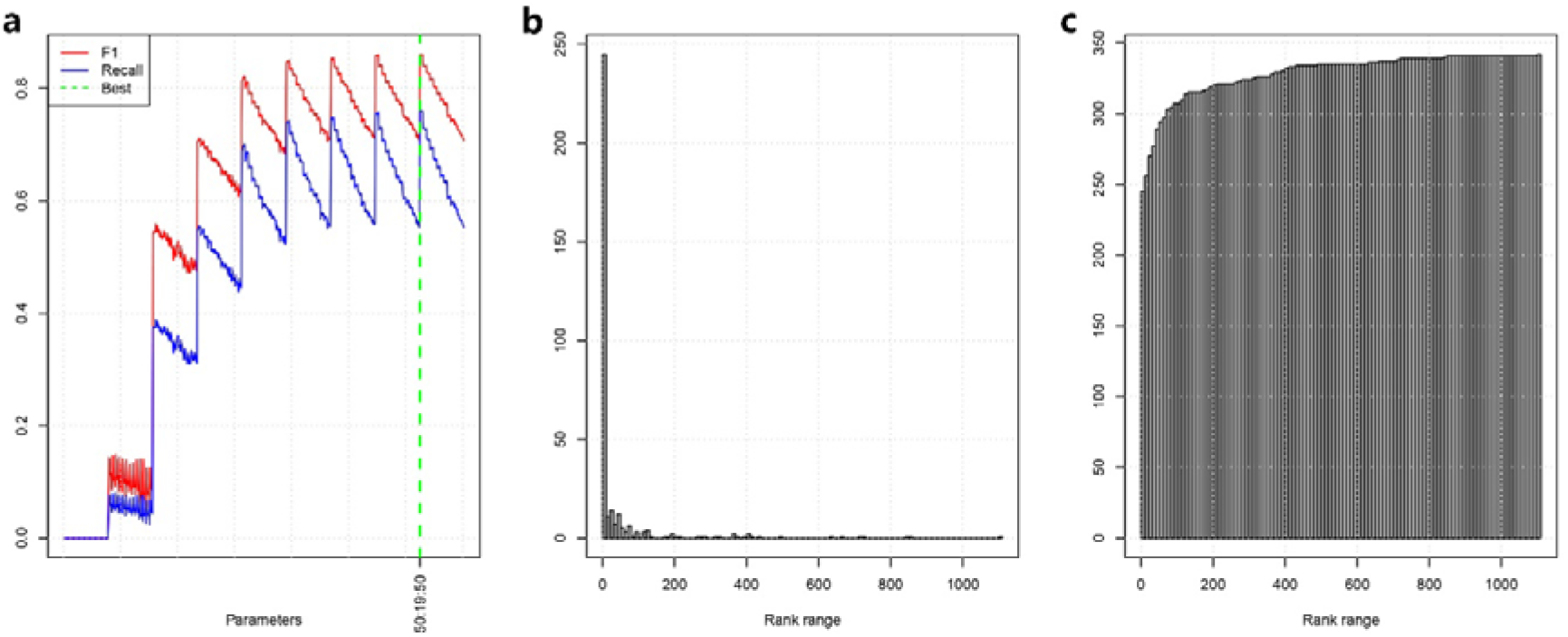
Performance in different parameter combinations. Y-axis represents the performance. X-axis represent the combinations of three parameters (n_estimators:max_depth:min_samples_split); 1b. rank distribution in cross-species validation. Y-axis is numbder and X-axis is rank; 1c. Cumulative distribution of the predictions.

To search HTH domain, we downloaded Pfam-A.hmm.gz (version 32) from Pfam database (21), then we extracted hmm containing HTH domains using hmmfetch in HMMER toolkit version 3.3 downloaded from http://hmmer.org/. We finally obtained 80 hmm files (Supplementary Data S1). To accelerate the HTH searching speed, we constructed a searching database using hmmpress program in HMMER toolkit. Finally, hmmscan in HMMER was employed to search Acr-associated domain under a cut-off of “-E 1e-5”.

### Model estimation

We leveraged precision, recall, accuracy and F1 score as measurements to estimate the capability of AcrDetector on the issue of screening Acrs. Precision can be calculated by the true positives (TP) divided by all predicted positives, which contains true positives and false positives (FP). Recall can reflect the ratio of detected positives in all positives. Therefore, precision and recall can reflect the capability to predict positive samples in two aspects. An F1 score is the harmonic mean of precision and recall, which is usually used to estimate model performance in an unbalanced dataset. The following four equations are used to define accuracy, precision, recall, and F1 score respectively. TN denotes true negatives, and FN denotes false negatives.

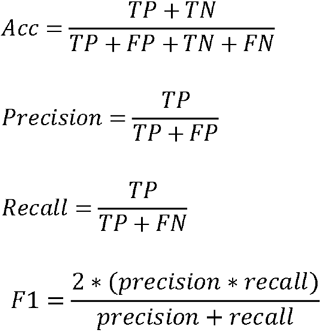

On the issue of determining Acrs, the performance of the classification model was estimated by multiround 5-fold cross-validations, independent dataset validation and unseen dataset test. Totally 3 estimation methods to evaluate AcrDetector’s performance.

### Classification model

We used “RandomForestClassifier” function in scikit-learn package (22) of python to execute the random forest model. To search the best parameters in the model, we utilized a greedy algorithm, grid-search cross-validation, to determine the best combination of three parameters: tree number in random forest tree (n_estimators), the maximum searching depth (max_depth) in a tree, and the minimum number for splitting a leaf node (min_samples_split). The search spaces were set as following: n_estimators from 50 to 100 with a step 10, max_depth from 3 to 20 using 2 as a step, and min_samples_split from 50 to 301 under 20 as an interval. We totally had 704 parameter combinations (Supplementary Data S2). Other parameters were default.

## RESULT

### Performance in 5-fold cross-validation

After traversing all the parameter combinations, we can achieve a best F1 score equal to 86.09% with 99.66% accuracy, 76.01% recall and 99.27% precision under n_estimators=50, max_depth=19 and min_samples_split=50 (Figure 1a). To test the robustness of AcrDetector, we performed other 30-round 5-fold cross-validation under different random states and the same parameters. We achieved a mean accuracy equal to 99.65% with variance 2.289e-09, a recall 75.84% with a variance 8.615e-06, a precision 99.24% with a variance 1.095e-06 and a F1 score 85.97% with a variance 3.106e-06.

### Validation in the whole genome scale

We simulated the practical situation in terms of searching Acrs. To do this, we need to access the probability for each prediction. A protein is more likely Acr if it is ranked in the top. We selected 159 species whose genome is assembled in the whole genome or chromosome level, regardless of other species in our benchmark, then performed prediction for each selected species after removing all sequences overlapped with the benchmark dataset in each prediction. Therefore, each prediction for each selected species can be regarded as an independent dataset test.

We assigned a rank to each Acr in each species according to the probability estimation. 71.43% Acrs are ranked in the top 10 (Figure 1b and 1c). Among the 159 selected species, only 42 species didn’t contain with any Acrs in its top ten predictions (Supplementary Data S3). In the top 15, 40 species didn’t contain with any Acrs (Supplementary Data S4). In all the positive predictions, there are 74.07% real Acrs (180/243) if we used probability of 0.5 as a cut-off, thus the precision is 74.07% which demonstrated our method can identify Acrs precisely.

### AcrDetector can identify three bona fide Acrs in unseen species

During our work additional Acrs have been published. AcrDetector was applied to those species (Supplementary Data S5 and https://github.com/RiversDong/AcrDetector/tree/master/examples). It accurately identified three bona fide novel Acrs (AcrIIA12, AcrIIA13b and AcrIIA19) in the top 15, but AcrRank didn’t identify them. The testing results on unseen species showed AcrDetector can identify true Acrs in unseen species. However, AcrRank can also identified an Acr, AcrIIA20, that did not determine by AcrDetector. Therefore, they can be jointly used to improve the precision. The results illustrated that our method can be used as a supplementary service of AcrRank, which is because AcrDetector can achieve better prediction than AcrRank for some species.

### Most of the Acrs are transferred into their host genome recently and AcrIE is expanded into closely related species after recently entering its host genome

Studies have shown that Acrs are usually located on MGEs. Here our result also illustrated Acrs tend to locate on genomic islands (GIs) and prophage segments (Figure 2a). Our *dev* index has shown that the codon *D* present obvious right skewed distribution, which illustrated very large codon usage bias between Acrs and non-Acrs (Figure 2b). Location on MGEs and large *dev* bias tell us that most of the Acrs are transferred into host genome recently, and Acrs do not adapt to the host genome codon. We investigated the Acrs distribution based on supplement data in reference (15). All the related species are group together in Figure 3. AcrIE was distributed as a single cluster, and they were not found in other species, which showed that AcrIE is expanded into closely related species after its recently enter its host genome, and then AcrIE becomes a young gene of the host species.

**Figure 2a.**
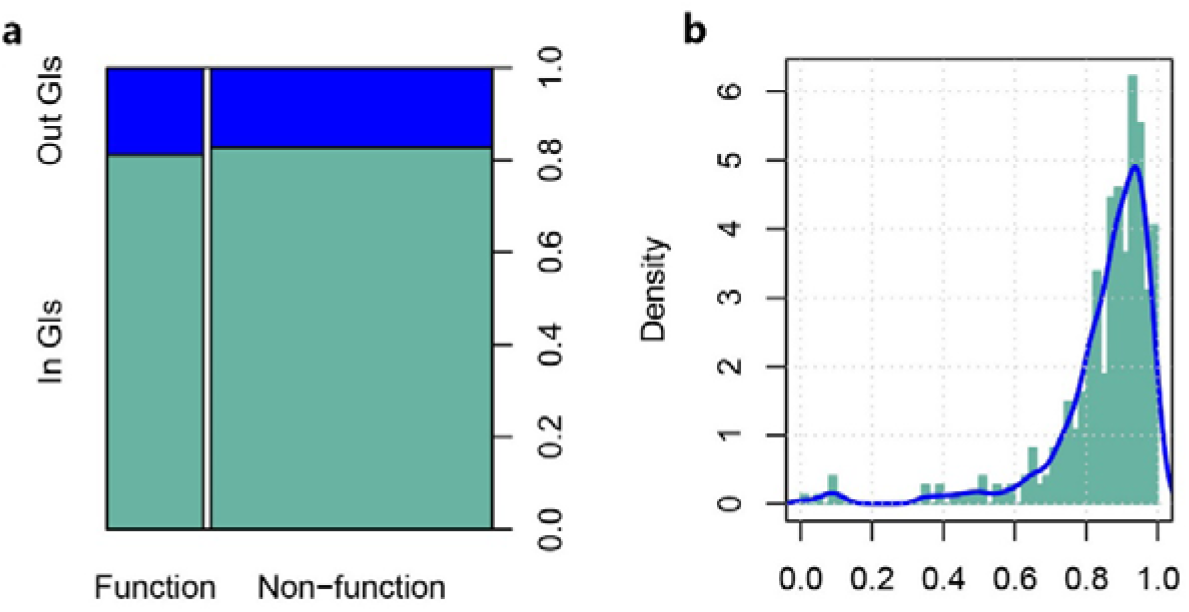
Location of *acr* genes (in or out of genome islands) and *acr* gene functions (hypothetical protein or non-hypothetical protein). Figure 2b *dev* comparison between *acr* and *non-acr*. Data that used to draw figure 2a is part of Acrs in the benchmark. Only species that assembled in whole-genome or chromosome level were used to draw figure 2b.

**Figure 3.**
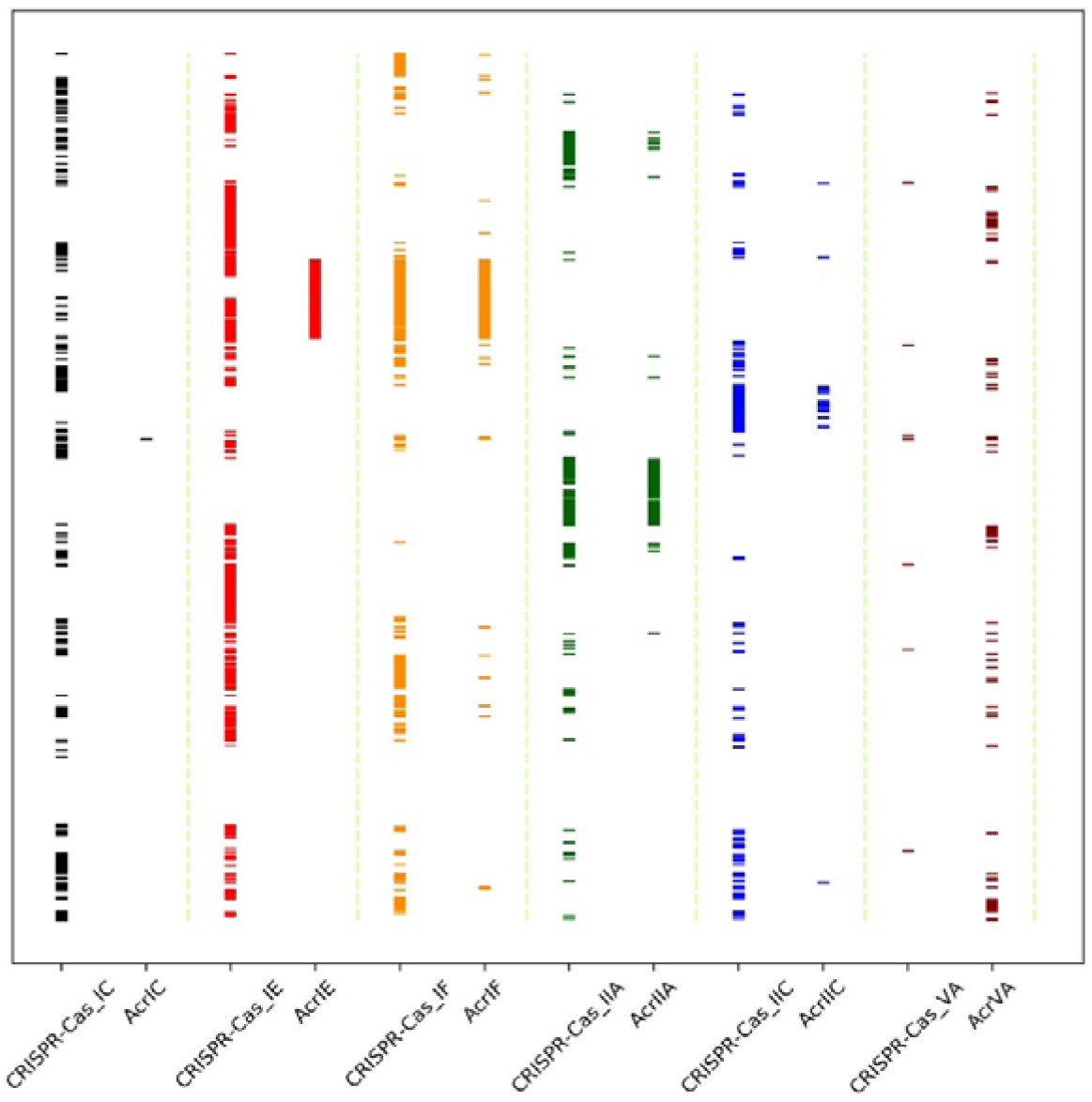
Distribution of Acrs and CRISPR-Cas systems. Data comes from Supplementary Data S2 of reference (15). Y-axis is species. All the species in Y-axis are ranked from alphabet A to Z, therefore species with similar names are clustered together.

## DISCUSSION

To be our knowledge, there are currently four tools to identify Acrs at the time of writing: AcrRank (http://acranker.pythonanywhere.com/), Acr catalog (http://acrcatalog.pythonanywhere.com/catalog/), AcrFinder (http://bcb.unl.edu/AcrFinder/index.php), and PaCRISPR (http://pacrispr.erc.monash.edu/index.jsp). AcrDetector, utilizes only 6 features and 50 decision trees in the random forest model to achieve a faster running time than the other four currently-available tools. The test on independent dataset and unseen species have shown the power of AcrDetector on the issue of determining Acrs. Additionally, only one feature, HTH, needs to perform domain search. Therefore, it’s easy to integrate it into other programs to perform large scale analysis. It’s difficult to pick out bona fide Acrs from the whole genome-scale because of the high sequence diversity. Therefore, depending on features derived from genomic background is promising. Five features among the six employed in AcrDetector are from the genome itself.

Some drawbacks of AcrDetector are presented here. Firstly, it can tell you which gene among your submitted genome is Acr, but it can’t tell you which CRISPR-Cas system it may inhibit. Users need to make a judgment by CIRSPR-Cas (sub)type in the prokaryotes. Some web servers and programs can be used to finish the judgment task such as CRISPRdisco (23), CRISPRCasFinder (24) CasLocusAnno (25) and CRISPRminer (26). Secondly, it only detects Acrs in prokaryotic genomes. For some Acrs in viruses, AcrDetector couldn’t search such genes. Thirdly, AcrDetector isn’t suitable for the genome with poor sequencing quality or low assembly level.

## SUPPLEMENTARY DATA

Supplementary Data S1: HTH domain used in this work

Supplementary Data S2: Performance in different parameter combinations

Supplementary Data S3: 159 selected species that used to cross-species (independent dataset) validation. It lists the rank details of Acrs.

Supplementary Data S4: 159 selected species that used to cross-species. The situation of the top 15 in our predictions.

Supplementary Data S5: predictions in unseen dataset

All scripts of the work can be downloaded from https://github.com/RiversDong/RiversDong.github.io/tree/master/ProjectOfAcrDetectorScript

## ACKNOWLEDGEMENT

We thank Dylan Sosa for his linguistic assistance during the preparation of this manuscript

